# Plant species vary in nectar peroxide, an antimicrobial defense tolerated by some bacteria and yeasts

**DOI:** 10.1101/2024.10.25.620320

**Authors:** Leta Landucci, Rachel L. Vannette

## Abstract

Floral nectar attracts pollinators but can also harbor microbes including plant pathogens. Nectar contains antimicrobial constituents including hydrogen peroxide, yet it is unclear how widespread nectar hydrogen peroxide might be among plant species or how effective against common nectar microbes. Here, we surveyed 45 flowering plant species distributed across 23 families and reviewed the literature for measurements of 13 other species to assess the field-realistic range of nectar hydrogen peroxide (Aim 1) and explored whether plant defense hormones induce nectar hydrogen peroxide upregulation (Aim 2). Further, we tested the hypotheses that variation in microbial tolerance to peroxide is predicted by microbe isolation environment (Aim 3), that increasing hydrogen peroxide in flowers alters microbial abundance and community assembly (Aim 4), and that microbial community context affects microbial tolerance to peroxide (Aim 5). We found that sampled plants contained nectar hydrogen peroxide ranging from 0 to 2940 µM, although half of the sampled species concentrations were less than 100 µM. While plant defensive hormones did not change the concentration of hydrogen peroxide in floral nectar, enzymatically upregulated concentrations in living flowers significantly reduced microbial growth. Together, our results suggest that nectar peroxide is a common but not pervasive antimicrobial defense among nectar-producing plants. In addition, nectar microbes vary in tolerance and detoxification ability, and co-growth can facilitate the survival and growth of less tolerant species, suggesting a key role for community dynamics in microbial colonization of nectar.

## Introduction

Many flowering plants offer energy-dense nectar to attract pollinators. While nectar is an essential attractant for pollinators, it also presents a liability (González-Teuber & Heil, 2009; Heil, 2011). In addition to transporting pollen, pollinators act as vectors for dispersal-limited bacteria and fungi (Antonovics, 2005; Harper et al., 2010; Herrera et al., 2009; Morris et al., 2020). Some are plant pathogens and enter plant via floral tissues (González-Teuber & Heil, 2009; McArt et al., 2014; Sasu et al., 2010) while other microbes primarily reside within floral nectar, a chemically diverse habitat rich in nutrients (Adler et al., 2021; Chappell & Fukami, 2018; Pozo et al., 2014). Microbial growth can modify nectar characteristics including pH, floral scent, sugar ratios and total concentration, amino acids, volume and temperature, which have been shown to alter pollinator foraging behavior (Aizenberg-Gershtein et al., 2015; Álvarez-Pérez et al., 2012a; de Vega et al., 2022; Herrera et al., 2009; Pozo et al., 2014; Rering et al., 2018; Schaeffer et al., 2017; Vannette & Fukami, 2018). However, many plant traits are hypothesized to protect nectar against potential larcenists including microbes (Adler, 2000; C. Carter & Thornburg, 2004). High sugar concentrations reduce the number of bumble bee-vectored yeast species able to survive within the nectar of *Helleborus foetidus* (Herrera et al., 2009). Undetermined chemical properties of nectar also reduce the growth of bacterial wilt in cucumber (Sasu et al., 2010).

These findings suggest potential adaptive value of nectar antimicrobial mechanisms (Adler, 2000; Herrera et al., 2009). However, much remains to be understood about the identity of and mechanisms by which floral nectar components are responsible for this microbial filtration, and whether strategies are conserved across phylogenetically diverse plant species. Floral nectar is a complex solution of metabolites and molecules in which carbohydrates, vitamins, lipids, amino acids, proteins, inorganic ions, and secondary compounds like alkaloids and phenolics are diverse and in some cases, abundant. (Adler, 2000; González-Teuber & Heil, 2009; Palmer-Young et al., 2019). Alkaloids, proteins and hydrogen peroxide inhibit microbial growth in nectar or nectar analogs (Aizenberg-Gershtein et al., 2015; Koch et al., 2019; Mueller et al., 2023; A. Schmitt et al., 2021; A. J. Schmitt et al., 2018a). These findings support the antimicrobial hypothesis, suggesting that specific nectar metabolites may be involved in adaptive defense and filtration of introduced microbes, significantly mediating microbial community composition and assembly in nectar (Adler, 2000; Harper et al., 2010; Herrera et al., 2009; Mueller et al., 2023; Pozo et al., 2014; Stevenson et al., 2017) yet how prevalent such mechanisms may be among plant species is poorly understood.

Hydrogen peroxide (H_2_O_2_) is a reactive oxygen species that generates free hydroxyl radicals (•HO) in the presence of metal ions. Hydrogen peroxide generation has been proposed to be a major mechanism of antimicrobial defense (C. Carter et al., 2007; C. Carter & Thornburg, 2004) in nectar, and more broadly across host-microbe systems (Allaoui et al., 2009; Miller et al., 2020; Smirnoff & Arnaud, 2019). Indeed, among potential antimicrobial compounds examined in laboratory studies, H_2_O_2_ had the broadest inhibitory effect across many microbial species (Mueller et al., 2023). In *Nicotian*a, nectarin proteins including superoxide dismutases and glucose oxidases drive a “nectar redox cycle” that produces hydrogen peroxide and also provides a mechanism for detoxifying the free radicals generated, allowing plants to avoid self-toxicity (C. Carter & Thornburg, 2000, 2004, 2004; González-Teuber & Heil, 2009; Harper et al., 2010). Hydrogen peroxide has been detected in the nectar of many species of *Nicotiana,* ranging between 23.4 and 4000 μM, but also at lower levels in *Cucurbita* and the legumes *Mucuna* and *Robinia* (Table 1) (Bezzi et al., 2010; C. Carter & Thornburg, 2000; Liu et al., 2013; Nocentini & Guarnieri, 2014; F. A. Silva et al., 2018a). However, whether antimicrobial levels of peroxide are also found broadly in the nectar of phylogenetically diverse plant species remains unknown. Hydrogen peroxide is also a common antimicrobial defense used by bees, found at high levels in bee-related habitats including the honey food stores of social Hymenoptera, produced via bee-secreted glucose oxidase proteins (Burgett, 1974).

**Table 1.**
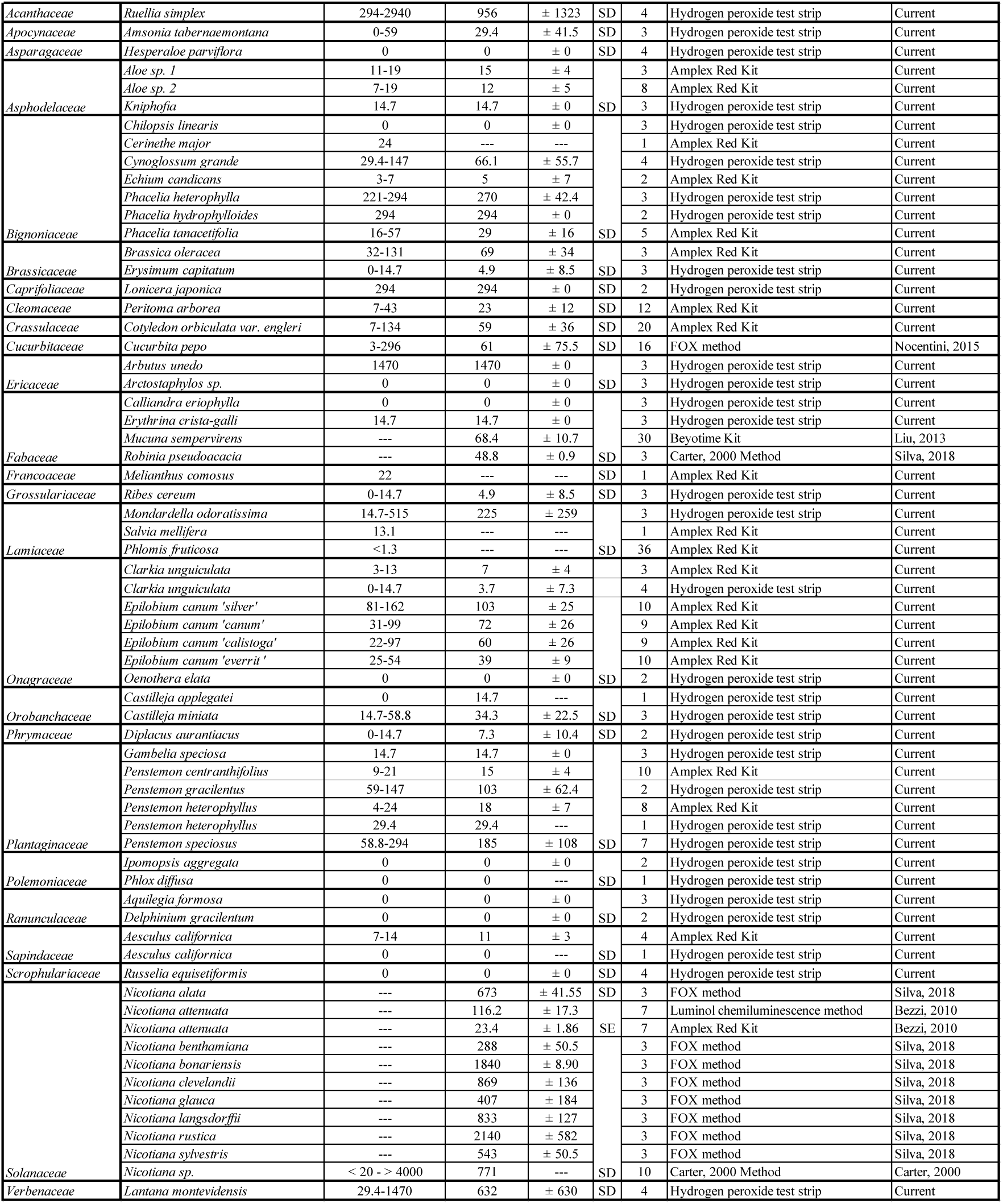
Hydrogen peroxide concentrations measured in nectar extracted from 58 different floral species spanning 25 families, both in this study, and reported in the literature. In the current study (45 species; 23 families), nectar samples from individual flowers were pooled to provide sufficient volume for either AmplexRed or colorimetric test strip hydrogen peroxide detection assays. SD and SE are abbreviations for standard deviation and standard error, respectively.

Concentrations of hydrogen peroxide at and above 2000 μM have been shown to reduce the growth of some common plant pathogens and yeasts and bacteria isolated from nectar, pollinators, and the environment (C. Carter et al., 2007; C. J. Carter & Thornburg, 2004; Mueller et al., 2023). However, some microbes may tolerate high hydrogen peroxide concentrations (Álvarez-Pérez et al., 2012; Herrera et al., 2009; Vannette et al., 2013), including the nectar-specialist yeast *Metschnikowia reukaufii*. Catalases defend against oxidative stress by catalyzing the decomposition of hydrogen peroxide, detoxifying the environment and potentially making it more favorable for microbial growth (de Vega et al., 2022; Vannette et al., 2013), including for the catalase producer and possibly co-occurring microbes (Mueller et al., 2023). This has yet to be evaluated in the complex and dynamic nectar environment of living flowers, or at concentrations of hydrogen peroxide lower than 2000 µM. Investigating whether lower concentrations of hydrogen peroxide, which may be more commonly found in nectar, are also antimicrobial, is necessary to assess the range of efficacy. Additionally, evaluating microbial taxa that are specialized to nectar and pollinator-associated environments and grow within a community, rather than as individual isolates, will be necessary to assess the range of conditions under which hydrogen peroxide may be an effective defense.

Here, we assess the generality of hydrogen peroxide as a potential antimicrobial compound in floral nectar, including its inducibility and variation in microbial tolerance and compositional response to peroxide. In Aim 1, we surveyed 45 plant species and compiled previous values for 13 additional species from the literature, spanning 25 families to measure the range of nectar peroxide concentration and examine if plant phylogeny predicts hydrogen peroxide concentration. In Aim 2, we assessed if plant defense hormones methyl salicylate and methyl jasmonate induce hydrogen peroxide upregulation in nectar. Salicylic acid plays a central role in priming plant defenses against potential microbial pathogens and we predicted that this could lead to increased nectar defenses as well, a possible mode of hydrogen peroxide regulation against microbes introduced into nectar. Next, we assessed microbial responses to field-relevant concentrations of hydrogen peroxide. We approached this question from multiple perspectives. In Aim 3, we tested how yeast identity or isolation source, including bee and flower-associated habitats, impacts tolerance to hydrogen peroxide using in-vitro assays. Next, in Aim 4 we examined how enzyme-increased hydrogen peroxide in the field shapes microbial community assembly in nectar. Finally, in Aim 5, we examined how individual vs. community context determines microbial growth and effects on peroxide concentration using lab assays. We predicted that differences we might observe could be mediated by microbial competition, or the benefits of co-growth, where the detoxification of hydrogen peroxide by more tolerant microbes could facilitate improved growth of less resilient microbes. Together, our results suggest that many but not all plant species maintain antimicrobial levels of peroxide, yet some common nectar and bee-associated microbes exhibit significant tolerance to high peroxide and can detoxify it, increasing growth of poorly-adapted microbes within a community.

## Methods

### Aim 1: Assessing natural levels of hydrogen peroxide in floral nectar

We sampled the floral nectar of 45 flowering plant species (Plantae: Angiospermae) (in addition to four cultivars of *Epilobium canum*) growing on the campus of University of California, Davis (38.540 °N, 121.756 °W, USA: California: Yolo County) and in the Sierra Nevada Mountain Range (Incline Lake, USA: Nevada: Washoe County) between March-July 2023 & 2024. We selected plant species that were in flower in spring/summer (2023 & 2024), produced nectar in volumes high enough to be extracted using glass microcapillary tubes (VWR International, Drummond Scientific Company, 50 μL) and have been previously studied with interest in microbial community inoculation (J. Cecala et al., 2024). To determine the concentration of nectar hydrogen peroxide we used one of two lab assays. For the first method, 5 μL of floral nectar was assayed using the colorometric Amplex^TM^ Red Hydrogen Peroxide/Peroxidase Assay Kit (ThermoFisher, Waltham, MA), and measured at an absorbance of 560 nm on a spectrophotometer microplate reader (model SYNERGY HTX) (See supplemental).

Alternatively, we used colorometric peroxide test strips with 0.5-2 μL of nectar (WaterWorks^TM^ Mid Range Peroxide Check, Industrial Test Systems Inc., Rock Hill, SC). Despite having lower precision, test strips exhibited a linear response to peroxide which the Amplex red kit did not, so the test strips gave us greater confidence in the range of measurement without the need to measure multiple dilution points (See Supplemental Methods). We note however, that this method was not necessarily specific to hydrogen peroxide, but any peroxides present and detected. We assessed the existing body of literature on hydrogen peroxide concentration in nectar, recording concentrations and detection methods used (Table 2).

**Table 2:**
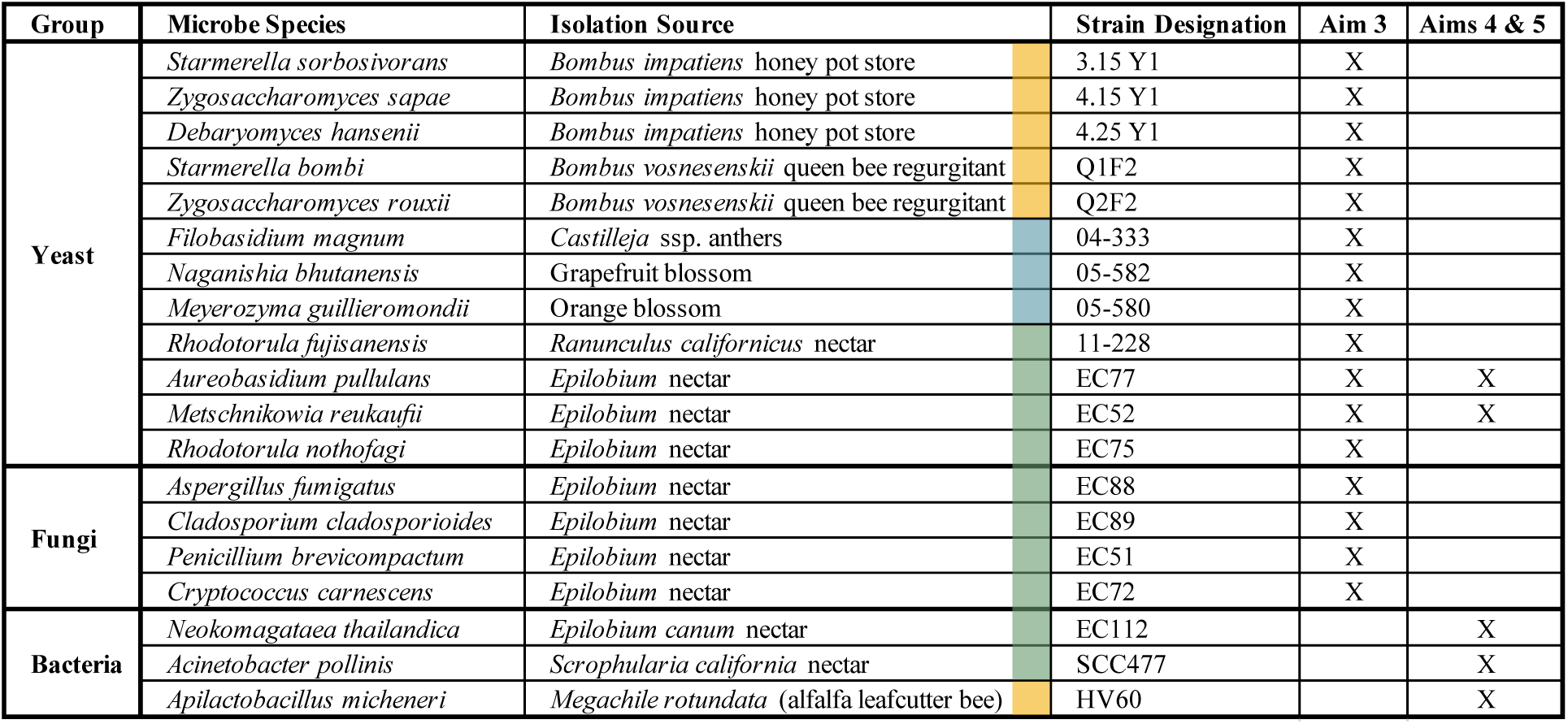
Fungi and bacteria studied in Aims 3, 4 and 5, with information about their growth form and isolation source.

### Aim 2: Does microbial defense plant hormone methyl salicylate induce upregulation of floral nectar hydrogen peroxide?

To explore whether the production of nectar hydrogen peroxide may be induced by common plant hormones in living flowers we tested the effects of methyl jasmonate and methyl salicylate. Methyl jasmonate and methyl salicylate upregulate defensive plant pathways associated with herbivory and microbial invasion, respectively (Bruinsma et al., 2008; Hernandez-Cumplido et al., 2016; Hoffmeister & Junker, 2017; Radhika et al., 2010). We were curious if either of these plant hormones increased concentrations of hydrogen peroxide in the nectar of long-blooming *Peritoma arborea* (bladderpod). We performed these manipulative field experiments on the campus of the University of California, Davis.

We selected two spatially separated (by 0.25 mi) patches of bladderpod. We treated recently opened flowers by spraying them with approximately 650 μL (350 mL spray bottle) of either a methyl jasmonate (1 mM in 2% EtOH), methyl salicylate (1 mM in 2% EtOH), or 2% ethanol (control) spray solution (Thaler et al., 2001). We held a container under the sprayed flower to catch any excess liquid and prevent the treatment from impacting other parts of the plant. Care was also taken to avoid flooding the flowers with liquid (and diluting the nectar solution) by positioning flowers so they faced downward when spray was applied, such that the lower parts of the petals, as well as the tips of the stigma and anthers received treatment. We then placed mesh bags over the flowers to prevent pollinator visitation and removal of floral nectar.

After 24 hours, we collected the individual treated flowers. Floral nectar was extracted using glass capillary tubes (VWR International, Drummond Scientific Company, 50 μL) and the nectar column’s length combined with a conversion factor based on the tube size was used to calculate the total volume of nectar extracted per flower. Peroxide concentrations were assessed using the Amplex^TM^ Red Kit (Thermo Fisher Scientific) using 5 μL of nectar, as previously described in Experiment 1 (also, see supplement).

### Aim 3: How does microbial identity and isolation source influence tolerance to hydrogen peroxide?

To test how microbial identity and isolation source impacts tolerance to hydrogen peroxide, we selected fungi isolated from floral petal tissue, queen bumble bee regurgitant, bumble bee honey pot stores, and various floral nectars. For each assay, each strain was freshly streaked from freezer stocks (Table 2) onto yeast media (YM) agar containing 0.1% v/v of 100 mg/mL chloramphenicol (antibiotic) in methanol solution to inhibit bacterial growth. After at least 72 h of growth on agar at 25 C, we created pure culture suspensions at 0.1 OD_600_ in sterile 20% sucrose solution (0.2 micron syringe filter). We prepared artificial nectar growth media (see Aim 3 recipe in supplemental methods) and sterilized it with a 0.2 micron sterile filter. We prepared hydrogen peroxide stock solutions through a set of serial dilutions into artificial nectar using stabilized 3% hydrogen peroxide (Thermo Scientific Chemicals) resulting in final in-plate concentrations of 0 μM, 100 μM, and 2000 μM.

Microbial growth was assayed by monitoring OD_600 nm_ within 96 well growth plates. Each well contained 120 μL of the artificial nectar growth media, 10 μL of appropriate hydrogen peroxide stock solution, and 20 μL of microbial culture suspension. Each treatment combination was replicated in triplicate, and negative controls lacking microbe inocula were run at each hydrogen peroxide concentration to ensure lack of microbial contamination. We incubated the microplates at room temperature in the dark and used a spectrophotometric microplate reader (model SYNERGY HTX, BioTek Gen5 Software) to measure optical density at 600 nm every 2 hours for 24 hours. Before each read the plates were shaken linearly for 30 seconds.

### Aim 4: How does hydrogen peroxide concentration shape microbial assembly in the nectar of living flowers?

To assess whether artificially elevated hydrogen peroxide in nectar affects the composition and growth of an inoculated microbial community, we performed manipulative field experiments with *Epilobium canum* (Greene) P.H.Raven (Myrtales: Onagraceae) on The University of California, Davis campus from June to August 2023. *Epilobium canum* was selected because of its harvestable nectar volume and floral phenology and longevity, typically 3-5 days.

We prepared synthetic microbial communities and enzyme solutions for floral inoculation experiments. We selected microbial species isolated from flowers or pollinators that were morphologically distinguishable when grown on media (Table 2: recipe in supplemental methods) (J. M. Cecala & Vannette, 2024). Each microbe was plated from freezer stock onto agar and incubated for at least 72 h of growth at 25℃. Then suspensions of each of the five species were adjusted to approximately 10^6^ cells using a hemocytometer and aliquots stored at - 80℃ (Cecala & Vannette, 2024). To create floral inocula the day of the field experiment, aliquots from all of the five species were thawed, combined, and brought up to ∼100µL with the freezer stock solution to yield an inoculum containing roughly 10^4^ c µL^-1^ of each of the five species, or equivalently 5 x 10^4^ total cells µL^-1^. The microbial inocula was kept on ice. Each of the species was proven to be successfully culturable from the individually aliquoted inocula on media plates prior to use in experiments. To manipulate the concentration of hydrogen peroxide in living flowers, we prepared a solution of the hydrogen peroxide generating enzyme, glucose oxidase (isolated from *Aspergillus niger*, Type VII, ≥1000,000 units/g, Sigma-Aldrich) at a concentration of 0.32 units of enzyme/μL in 40% sugar solution (80% glucose: 20% sucrose), immediately prior to beginning the experiment. This solution, as well as a boiled (inactive, boiled for 2 hr) glucose oxidase solution at an equivalent concentration to act as a negative control were kept on ice. This strategy was advantageous because (1) the nectar redox cycle nectarin protein NEC5 identified in *Nicotiana* is a glucose oxidase and produces hydrogen peroxide via the same mechanism (C. Carter & Thornburg, 2004), (2) hydrogen peroxide can rapidly degrade, so this enzymatic method allows for a renewal of hydrogen peroxide through the activity of the GOX, and (3) a previous study showed success of this method in living *Nicotiana* nectar (Bezzi et al., 2010).

The day prior to our field experiment we covered closed flowers of *Epilobium canum* in two discrete patches with mesh bags to exclude pollinator visitation and microbial deposition (Francis et al 2023). Once all microbial inocula and enzyme solutions were prepared fresh each day, we identified bagged flowers that had opened and could be used in our experiment. For the first experimental treatment, we inoculated flowers with 10 μL (3.2 units) of active glucose oxidase enzyme and 1 μL of microbial inocula (n = 10, split between two clumps). Control flowers were inoculated with 10 μL of 40% sucrose solution and 1 μL of microbial inocula while additional flowers were inoculated with 10 μL boiled glucose oxidase solution and 1 μL of microbial inocula. Solutions were delivered into flowers using a micropipette and sterilized 10 μL tip. The inoculated flowers were marked with permanent marker, tagged, and mesh bags replaced. Each treatment had 10 replicates per trial, and two separate trials were conducted. The prepared microbial inoculum, the active glucose oxidase, boiled GOX, and control solution (sugar substrate) were all kept on ice while out in the field.

After 24 hours, flowers were harvested and brought back to the laboratory where microbial growth and composition were assessed. Occasionally, we would find that thrips and ants had made it inside of some bagged flowers and their presence noted. Nectar was extracted in a hood using glass capillary tubes, total nectar volume of each flower quantified (VWR International, Drummond Scientific Company, 50 μL), and diluted for plating. We also noted any bleached corolla or pistil tissue, likely caused by oxidative tissue damage from reactive oxygen derivatives of hydrogen peroxide. For floral nectar samples with greater than 5 μL extracted volume, we added 5 μL of nectar into 45 μL of DPBS buffer. For extracted nectar samples of less than 5 μL, 50 μL DPBS buffer was added to the full amount of nectar sample. We vortexed and centrifuged the diluted samples, then plated 15 μL of each sample on YM, TSA, and MRS plates and spread using sterile glass beads. Plates were incubated at 28 ℃ for 7 days. CFUs were identified by microbial taxa based on morphological features and counted for each plate (Cecala & Vannette, 2024). For plates where the CFUs were too numerous/dense to count with confidence, we overlayed a transparent grid and tallied CFU subsamples in at least six 0.1 or 1.0 cm^2^ squares, which we used to estimate the total CFUs per plate for a given microbe. If CFU’s were too dense for even this method, having formed a lawn, we assigned it the highest countable density from another plate.

Using 5 μL undiluted aliquots of extracted nectar above, we quantified hydrogen peroxide concentrations with peroxide test strips (WaterWorks^TM^ Mid Range Peroxide Check, Industrial Test Systems Inc., Rock Hill, SC) to validate enzyme activity and examine if microbial growth affected peroxide concentration.

### Aim 5: How does community vs individual growth context impact microbial growth?

To explore whether co-growth of microbes influences microbial tolerance to hydrogen peroxide, we compared the growth of microbe isolates individually and in communities at different concentrations of hydrogen peroxide, in sterile artificial nectar solution to mimic floral nectar conditions (see Aim 5 recipe in Supplemental Materials). Using the same microbial isolates also used in the previous field experiment (Table 1), we created a community suspension of all five taxa as well as suspensions for each individual taxa. To create a community inoculum the day of the in vitro lab experiment, we combined aliquots from each of the five species as described previously. To make microbial isolate suspensions, an aliquot from a given species containing roughly 10^4^ c µL^-1^ was brought up to a volume of 100 μL with freezer stock solution. We prepared hydrogen peroxide stock solutions through a set of serial dilutions into the artificial nectar media with final concentrations of 0 μM, 10, 50 μM, 100 μM, and 2000 μM.

For each replicate, we added 39 μL of artificial nectar, 10 μL of hydrogen peroxide treatment and 1 μL of microbial suspension for a total volume of 50 μL in PCR tubes, which we selected based on nectar volumes often extracted from *Epilobium canum* during prior field trials. Each treatment combination--six different microbial inocula (community of all five microbes and each microbe individually) x five hydrogen peroxide treatment concentrations-- was performed in triplicate. We vortexed and centrifuged the tubes and then incubated at 28℃ for 24 hours. After the 24 hours we assessed microbial growth as described in Aim 4 above. We also determined the concentration of hydrogen peroxide using 5 μL of each solution applied to peroxide colorimetric test strips. By comparing control, individual and community samples we assessed how microbial growth and community membership altered hydrogen peroxide concentration.

### Statistical Analysis

We conducted all statistical analyses in R (R Core Team, 2021) and used ggplot (Wickham, 2016). In Aim 1, to estimate plant phylogenetic relationships among the sampled plant species, we used the function ‘phylo.maker’ in package *V.PhyloMaker2* (Jin & Qian, 2022) using the reference vascular plant mega-phylogeny GBOTB.extended.TPL. Using this tree and the function ‘multiPhylosignal’ in the package *picante* (Kembel et al., 2010) we tested for a phylogenetic signal of nectar hydrogen peroxide concentration.

To test whether defensive growth hormones altered concentrations of hydrogen peroxide in *Peritoma arborea* nectar or nectar volume (Aim 2), we used one-way ANOVA in the package *car*. We obtained F- and P values and Kenward-Roger degrees of freedom. For Aim 3, we tested whether microbial growth differed among concentrations of hydrogen peroxide according to microbial identity or isolation source and their interaction using type III ANOVA using the drop1 function. We performed post-hoc analysis using the ‘TukeyHSD’ function to assess the source of variation in the predictor variable, microbial isolation source. For Aim 4, we compared how enzyme treatments influenced floral nectar hydrogen peroxide concentration, total microbial community composition, and growth of individual species using separate ANOVAs. To evaluate which enzyme treatments contributed to differences observed in total microbial community composition and individual microbe growth, we performed a post-hoc Tukey analysis.

In Aim 5, we examined the impact of community vs isolate growth context, hydrogen peroxide concentration and their interaction on microbial growth using III ANOVAs, with separate models for each tested microbe. Additionally, we examined if measured peroxide concentration varied with microbial growth context, starting hydrogen peroxide treatment concentration and their interaction using ANOVA. To evaluate which microbe taxa contributed to differences observed in the final environmental hydrogen peroxide concentration, we performed a post-hoc analysis using ‘TukeyHSD’. Finally, to test if microbial community composition (as Bray-Curtis dissimilarity) that formed after 24 hours differed among hydrogen peroxide treatments, we used function ‘adonis’ in package *vegan* (Oksanen et al., 2007)to perform a multivariate analysis of variance (PERMANOVA). Microbial community composition was visualized using non-metric multidimensional scaling (NMDS) ordination.

## Results

### Aim 1: Plant species vary in hydrogen peroxide concentration in floral nectar

Across 45 species sampled here, peroxide concentrations in floral nectar ranged from 0 to 2940 μM across species (Table 1; Figure 1). *Arbutus unedo* nectar averaged the highest sampled hydrogen peroxide concentration, measuring 1470 μM ± 0 (SD), and was followed by *Ruellia simplex* (956 μM ± 1323) and *Lantana montevidensis* (632 μM ± 630). These values were well within the range of hydrogen peroxide measurements in the literature for *Nicotiana,* which as a genus, averaged 923.3 μM ± 636. Within the entire dataset (Table 1; 58 species; 25 families) only 29% of the sampled plant species had hydrogen peroxide concentrations between 100 and 2000 μM and only 2% of species had concentrations >2000 μM. Meanwhile, concentrations between >0-100 μM were found in 52% of the species. About 17% had no detectable levels of hydrogen peroxide. Among the surveyed plant species, phylogenetic relatedness did not predict hydrogen peroxide concentration (K = 0.15, PIC.variance.P = 0.56) (Figure 1). This result likely reflects the large variation observed within some clades, however we note that some plant clades exhibit consistently high (*Nicotiana*) or low peroxide concentrations given the plant species included here (e.g. Monocots; Brassicaceae).

**Figure 1.**
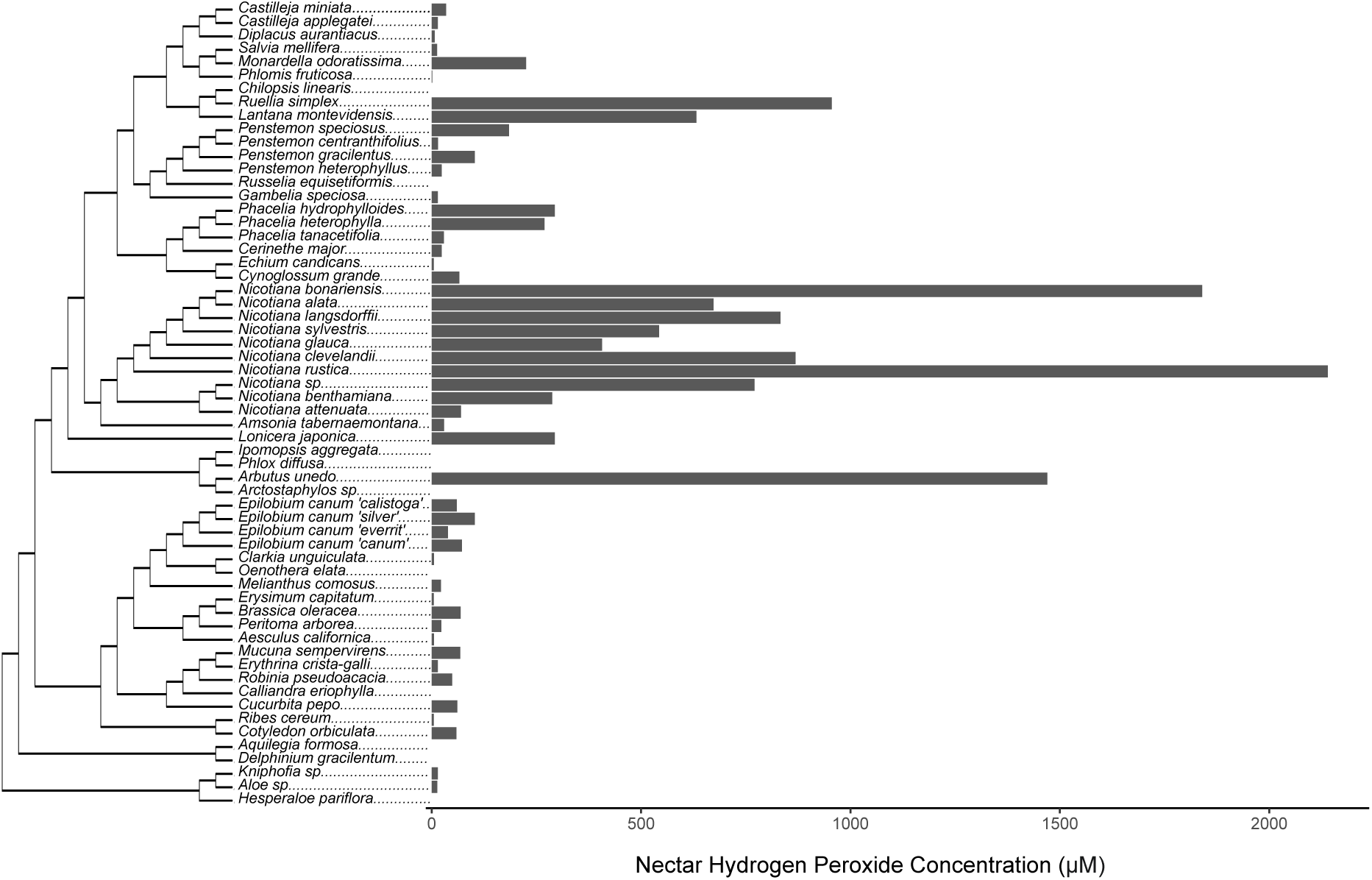
Plant species varied in peroxide concentration in floral nectar but no significant phylogenetic signal was detected among flowering species surveyed (K=0.15, PIC.variance.P=0.56). Plant species include newly sampled and literature values (Table 1).

### Aim 2: Plant defense hormones do not elicit changes in nectar peroxide in *Peritoma arborea*

We investigated whether nectar hydrogen peroxide may be an induced defensive response to plant hormones methyl salicylate and methyl jasmonate. We found that neither methyl jasmonate nor methyl salicylate significantly affected concentration of hydrogen peroxide in *Peritoma arborea* nectar (F_2,77_ = 0.25, *p* = 0.78) (Figure 2A) and did not affect nectar volume (F_2,77_ = 1.17, *p* = 0.32) (Figure 2B).

**Figure 2:**
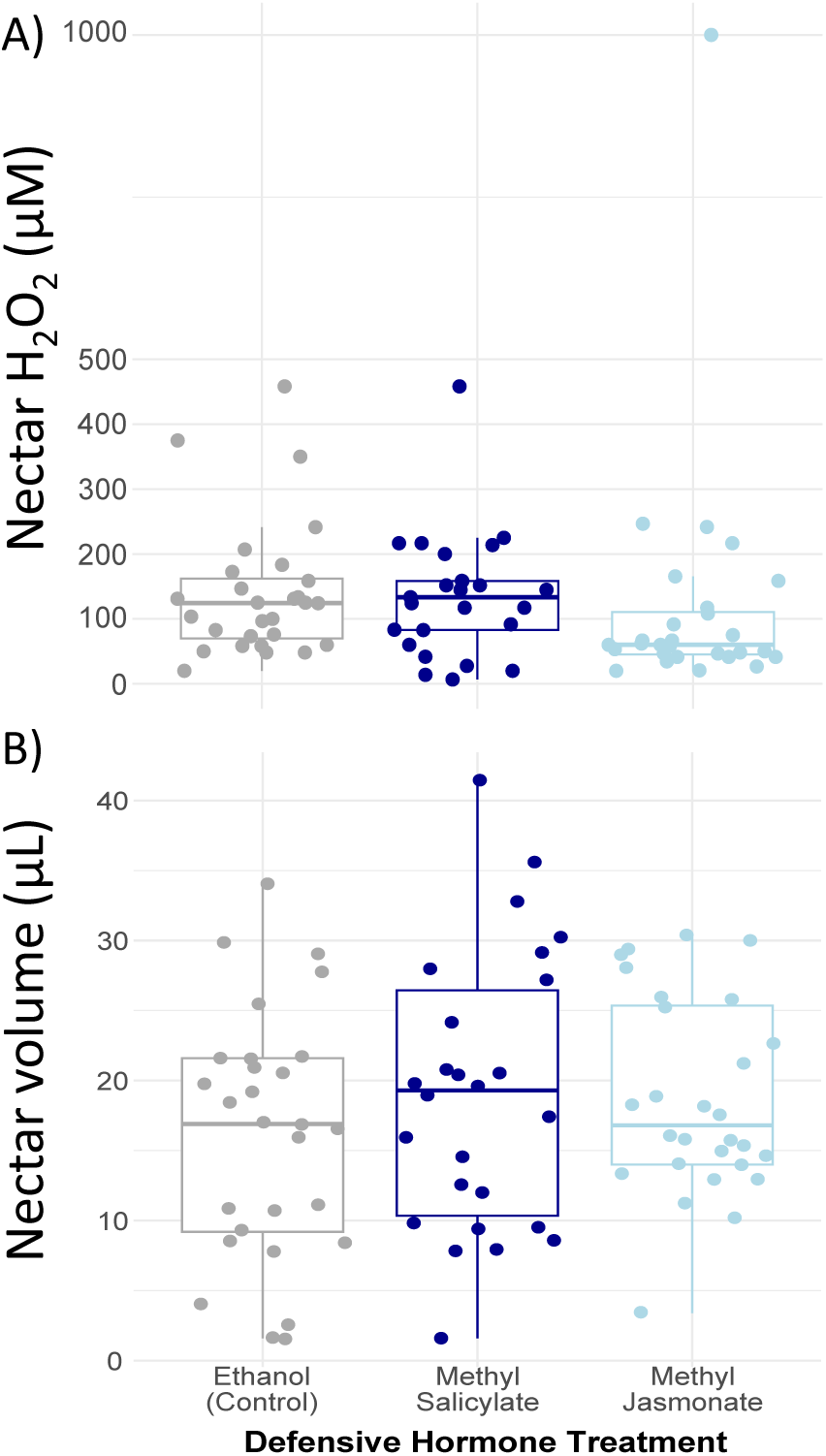
No effect of plant hormones methyl salicylate or methyl jasmonate on a) hydrogen peroxide in nectar of *Peritoma arborea* (F_2,77_ = 0.25, p = 0.78), nor on B) nectar volume (F_2,77_ = 1.17, p = 0.32). Hydrogen peroxide concentration was measured with the AmplexRed detection assay. Boxplot center line indicates the median and outer lines indicate upper and lower quartiles, with whiskers including 95% CI. N=26-28 flowers/treatment.

### Aim 3: Variation in microbial tolerance to peroxide is not predicted by isolation source

We assessed peroxide tolerance of fungal taxa isolated from various sources (floral petal tissue, queen bumble bee regurgitant, bumble bee honey pot stores, or floral nectar, Table 2). Microbial taxa exhibited variation in tolerance to hydrogen peroxide concentration (Figure 3A; Peroxide x species F_30,239_ = 7.54, *p* < 2e^-16^), but in contrast to our predictions, microbial isolation source was not a significant predictor of growth responses to peroxide (Fig 3B; Source x H_2_O_2_ concentration F_4, 278_ = 1.26, *p* = 0.29). Many microbes, including *Debaryomyces hansenii* and *Aureobasidium pullulans,* experienced extreme reduction in optical density (OD) when grown at 2000 μM H_2_O_2_, with *D. hansenii* OD reduced by 10 fold and *A. pullulans* by 8.75 fold. In contrast, yeasts from various environments showed limited growth reductions to all assayed hydrogen peroxide concentrations including 2000 µM (Fig 3).

**Figure 3:**
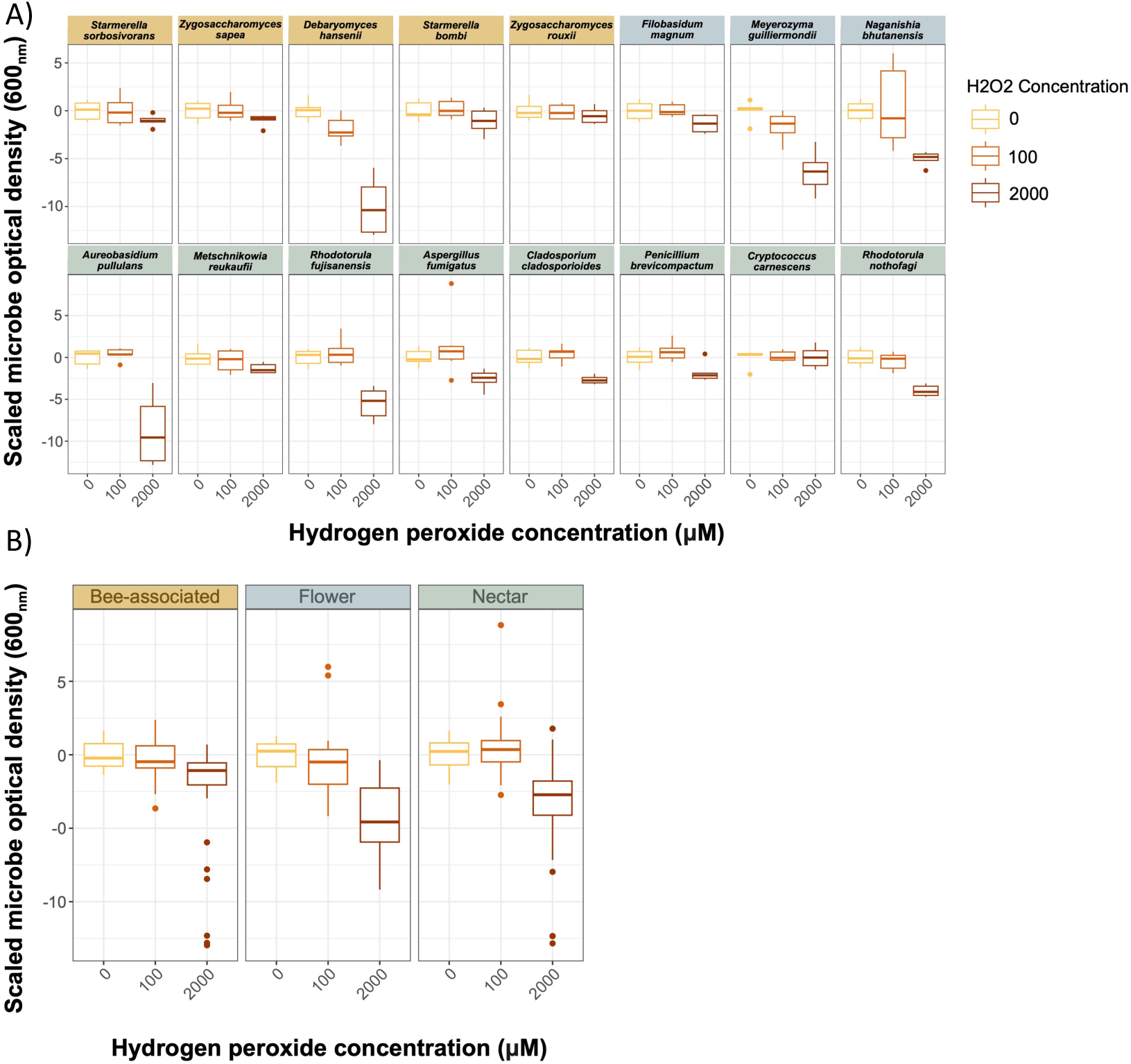
Increasing hydrogen peroxide concentration reduces yeast growth depending on **A)** microbial species identity (Peroxide x species F_30,239_ = 7.54, *p* < 2e^-16^) but does not vary with **B)** isolation source (Source x H_2_O_2_ concentration F_4, 278_ = 1.26, *p* = 0.29). N= 6 for each microbe-concentration. Boxplot center line indicates the median and outer lines indicate upper and lower quartiles, with whiskers including 95% CI.

### Aim 4: Elevated peroxide in nectar reduces microbial abundance and diversity in situ

We manipulated the concentration of hydrogen peroxide in living *Epilobium canum* flowers with glucose oxidase enzyme and measured its impact on the growth of five microbial taxa (including two fungi used in Aim 3). Enzyme addition significantly increased peroxide levels beyond our quantification threshold, exceeding 11760 μM (F_2,47_ = 153.02, *p* < 2e^-16^) while flowers inoculated with inactive boiled glucose oxidase averaged 4 µM in hydrogen peroxide concentration and the control sugar solution measured at 0 μM. Addition of the active glucose oxidase enzyme significantly reduced microbial growth overall (Fig 4B), suppressing the growth of all microbes that could be recovered from nectar (treatment x microbe F_8,352_ = 9.31, *p* = 1.15^e-11^, Figure 4C). Only *M. reukaufii*, a nectar specialist yeast, was recovered from a subset of flowers treated with glucose oxidase (n = 7 flowers, ∼23% of those inoculated), suggesting its ability to withstand extremely high peroxide levels. Interestingly, inactivated enzyme addition enhanced the growth of bee-associated bacterium *A. micheneri* (Figure 4C). This bacteria was not detected in the sucrose treatment or in the presence of the active enzyme, suggesting microbial utilization of nutrients from the degraded enzyme allowed it to persist but this was not sufficient to tolerate high peroxide concentration.

**Figure 4.**
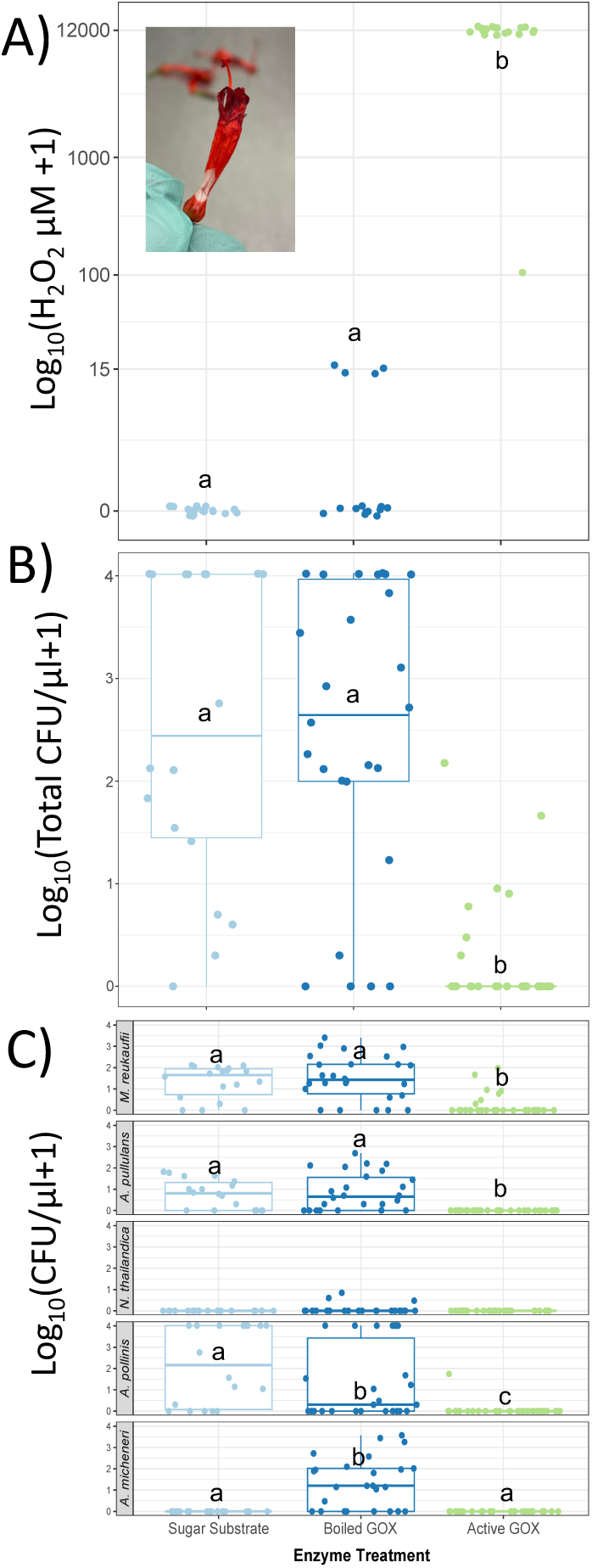
Enzyme addition to flowers alters **A)** hydrogen peroxide concentration in *Epilobium canum* floral nectar via inoculation with glucose oxidase enzyme (GOX) and sugar substrate (F_2,47_ = 153.02, *p* < 2e^-16^). Peroxide concentration was measured using peroxide test strips.

Boiled GOX and sugar substrate treatments serve as negative controls. Samples were collected over two days from two spatially distinct plant clumps (not the same flowers used in figures B & C). N/treatment = 11-20. Letters indicate statistical difference between treatment groups. Photo inset shows bleached floral tissue from GOX treatment. **B**) The effect of glucose oxidase on total microbial community density (CFU/uL nectar), determined by counting the number of colony forming units (CFUs) cultured on media plates after extracting floral nectar. The microbe community consists of five microbes, two fungi and three bacteria. **C)** The impact of glucose oxidase on the CFUs of individual microbe taxa within the inocula (treatment x microbe F_8,352_ = 9.31, *p* = 1.15^e-11^), as well as the influence of microbial identity on growth outcomes (P=0.013). Samples for B and C were collected over two days and from two spatially distinct clumps of *Epilobium*. N = 19-20 for GOX and boiled GOX samples, and N = 8 for all samples treated with sugar substrate. Boxplot center line indicates the median and outer lines indicate upper and lower quartiles, with whiskers including 95% CI.

### Aim 5: Community context mediates microbial response to peroxide concentrations and peroxide degradation

We compared growth of microbes in community to microbes grown alone at various hydrogen peroxide concentrations in a laboratory in vitro setting (Figure 5A). For most microbes, community context mediated growth, and for two microbes, community context mediated effects of peroxide on growth (*M. reukaufii* context x peroxide: F_1,116_ = 5.44, *p* = 0.021, *A. micheneri* context x peroxide: F_1,116_ = 7.75, *p* = 0.0062). Yet the type and direction of microbial responses to context and peroxide varied. For example, *M. reukaufii* grew better individually than in community (F_1,116_ = 149, *p* = <2^e-16^) and increased growth with greater peroxide concentration (F_1,116_ = 7.4, *p* = 0.0075). Conversely, *A. michener*i grew significantly better when in a community and its response to high peroxide depended on community context (F_1,116_ = 12.7, *p* < 0.00052). In contrast, *A. pullulans* and *N. thailandica* only grew individually (F_1,116_ = 49, *p* = 1.76^e-10^; F_1,116_ = 12.6, *p* = 0.00055) and did not respond significantly to variation in peroxide.

**Figure 5.**
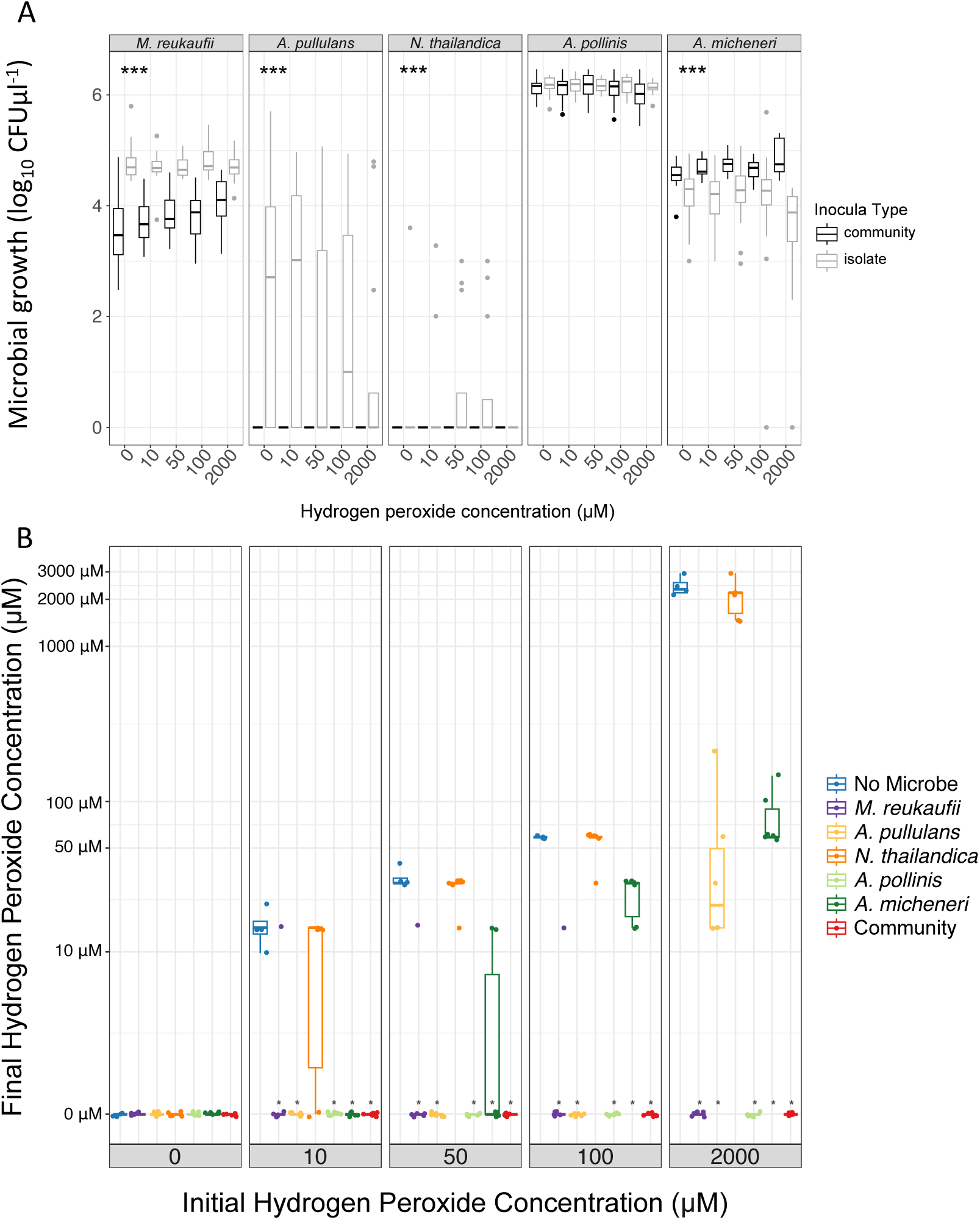
A) Hydrogen peroxide differentially affects microbial growth depending on community context (co-growth as black and individuals as gray) on microbial growth (log10 transformed CFU count). N=12 for each microbe-community complexity-peroxide combination. N = 4 for negative controls for each treatment. Microbes with * grew significantly different between community growth contexts. Boxplot center line indicates the median and outer lines indicate upper and lower quartiles, with whiskers including 95% confidence intervals. **B)** Microbe growth affects final hydrogen peroxide concentration but depends on species and community context after a 24 hour incubation period. N = 6 for each sample/treatment type. N = 4 for the no microbe controls.

Meanwhile, only a single microbe, *A. pollinis,* was unaffected by context (F_1,116_=1.1, P=0.27) but was affected by peroxide concentration (F_1,116_ = 3.97, *p* = 0.049), with increasing peroxide reducing its growth. We found no significant effect of hydrogen peroxide concentration on microbial composition using lab assays (PerMANOVA: F_4,58_ = 0.75, *p* = 0.63).

Some but not all microbes significantly reduced the concentration of peroxide in analog nectar growth media relative to the control solutions (no microbe) (F_24,165_ = 102.39, *p* < 2^e-16^) which was increasingly evident with increasing initial hydrogen peroxide concentration (Figure 5B). For example, *M. reukaufii* and *A. pollinis* eliminated detectable levels of peroxide in all solutions after growing in artificial nectar for 24 hours. In contrast, *A. pullulans* and *A. micheneri* varied in their effects on hydrogen peroxide in solution, reducing concentrations at low initial concentrations but not at higher concentrations. The bacteria *N. thailandica* did not significantly reduce hydrogen peroxide at any treatment concentration despite exhibiting growth (Fig 5A).

When all 5 microbes were grown together as a community, hydrogen peroxide concentrations were entirely quenched at all treatment levels.

## Discussion

Together, our results suggest that nectar hydrogen peroxide is a common but not pervasive antimicrobial defense among nectar-producing plants. In addition, the microbes that colonize nectar vary in tolerance to and detoxification ability of peroxide, suggesting key adaptations to growth in an environment where oxidative stress is common. Finally, we document how co-growth can in some cases, facilitate the survival and growth of less tolerant species, suggesting a key role for community dynamics in microbial colonization of nectar and tolerance to its defenses.

Across plant species, we documented a wide range of peroxide concentrations, and strongly antimicrobial levels of peroxides (>1000 µM) in at least three plant orders (Fig 1; Lamiales, Solanales, Ericales). We detected no significant phylogenetic signal in peroxide concentration found in nectar, and noted variation among species within families. In contrast to this more general pattern, however, species in the genus *Nicotiana* hosted consistently high hydrogen peroxide values. This clade may exhibit unique nectar secretion mechanisms or biochemistry (Bezzi et al., 2010; Hillwig et al., 2010). We also note that all measurements within this clade were taken from previous studies and most were quantified with different methods than those employed here. Both within and outside the Solanales, however, our sampling suggests that many plant species do not present nectar with high peroxide concentrations (Table 1). In addition, we found no evidence that peroxide is inducible with either methyl salicylate or methyl jasmonate in bladderpod (*Peritoma arborea*). However, we note that plant species vary in plastic responses to defense hormones (Karban, 2011; Valladares et al., 2007) so this does not preclude induced nectar defense in other plant species or contexts. Yet the variation in peroxide concentrations among plant species suggests that peroxide is not necessarily a universal antimicrobial defense in nectar (Hillwig et al., 2010). We suggest a few explanations for such variation. First, nectar protein content and composition is variable among plant clades (F. A. Silva et al., 2018b; FredyA. Silva et al., 2020), so the biochemical pathways that produce strong oxidative chemistry in nectar may vary in presence or activity. Yet hydrogen peroxide and enzymes involved in its production are common among plant tissues and across species, so it would be surprising if biochemical constraints limit its frequency among plant species. Second, it may be that the cost of maintaining strong oxidative chemistry or quenching in nectar is high.

For example, we noted ‘bleaching’ damage to tissues (Figure 3A, inset) when we artificially increased hydrogen peroxide concentrations. Although some plant species including *Nicotiana* produce antioxidants to reduce self-damage, other species may experience significant cost in maintaining the chemistry for tolerating or quenching excess free radicals should they generate high peroxide levels in the first place. Instead, plants may evolve to tolerate microbial growth or invest in alternative antimicrobial strategies. Additional enzymatic defensive strategies in nectar have been characterized, including the activity of RNases (Hillwig et al., 2010), lipases (Kram et al., 2008) or antifungal lipid transfer proteins (A. J. Schmitt et al., 2018b). We also note that in addition to its proposed antimicrobial functions, peroxide generation can serve various physiological functions within plants (Smirnoff & Arnaud 2018) including some co-opted for novel functions including pigment generation in nectar (Magner et al., 2023).

While high peroxide and ROS production may suppress the growth of some microbes, we show that yeasts and bacteria from multiple isolation environments vary in tolerance, with some species exhibiting minimal reduction in growth at even 2000 µM H_2_O_2_. Detoxification mechanisms used by bacteria and fungi include peroxidases and catalases to degrade H_2_O_2_, which microbes also generate as a byproduct of metabolism. Indeed in nectar, most bacteria and fungi are catalase positive (Álvarez-Pérez et al., 2012b; Jacquemyn et al., 2013) yet even such bacterial and fungal species differ in the extent to which they are able to degrade peroxide and repair oxidative damage (Sen & Imlay, 2021). Our results confirm that such variation in tolerance and detoxification exists even in nectar and bee-associated microbes (Figs 2, 5).

Evidence presented here also supports the hypothesis that community context affects microbial assembly and composition, mediated via interactions with chemical stressors (Álvarez-Pérez et al., 2019). For example, effective peroxide detoxifiers (here, *M. reukaufii*) experienced competitive release at high peroxide concentrations. At the same time, these species may buffer competitor survival within these stressful habitats (Ma & Eaton, 1992) via quenching or other peroxide reduction mechanisms, as has been documented in other systems (Piel et al., 2021). It may be that previous reports of potential facilitation among nectar microbes (Alvarez-Perez & Herrera, 2013; Francis et al., 2023) have occurred in plants with high peroxide concentrations or involve similar detoxification mechanisms. Indeed, experiments presented here and in previous work (J. Cecala et al., 2024; Mueller et al., 2023) suggest that *Metschnikowia reukuafii* is an effective detoxifier of peroxides and can survive very high concentrations (>10,000 µM) of peroxide, and can facilitate microbial survival in high peroxide environments. We also note that in nectar environments, plant secreted enzymes continually produce peroxides while lab assays performed here used stabilized peroxides that were not regenerated. Our observation of microbial detoxification of peroxides occurred in lab conditions, so remains to be tested using in-plant assays. Yet we observed *M. reukaufii* tolerance to high peroxide concentrations in both lab and in situ conditions, supporting consistent responses. In any case, it is possible that co-inoculation or priority effects, or early arrival by some species, may determine the survival of species with lower peroxide tolerance (Álvarez-Pérez et al., 2019; Chappell & Fukami, 2018).

Our experiments raise further questions regarding the ecological costs and consequences of peroxide production in nectar. Previous work suggests that honey bees can detect, and in some cases avoid peroxide-supplemented nectar solutions (Bartlett et al., 2022; Bezzi et al., 2010), and that although bees consume high concentrations of bee-produced peroxide in honey, its detoxification may be costly (Korayem et al., 2012). It may be that high levels of peroxide in nectar incur ecological costs of reduced pollination services or altered pollinator composition, limiting the evolution of this defense among plant species, yet this hypothesis requires further examination. In addition, although peroxide generation biochemistry within *Nicotiana* is well-characterized, a variety of ROS generation reaction mechanisms are possible (Becker et al., 1990), including generation by secondary compounds in nectar (Magner et al., 2024). In conclusion, our experiments support the hypothesis that peroxides can act as an antimicrobial defense against many microbes in nectar yet highlights variation in microbial tolerance to such defenses, particularly in a community context.

### Statement of authorship

LL and RLV conceived of and designed all studies. Data collection was performed by LL with help from RLV. LL performed analyses and wrote the first draft of the manuscript with substantial input and revisions from RLV throughout the writing process. All authors read and approved of the final manuscript.

### Data availability

Raw data and code are provided in the supplementary materials for review and will be made available on Dryad or FigShare upon acceptance.

## Acknowledgments

We thank all Vannette lab members including Jacob Cecala for coding assistance, Danielle Rutkowski, Alexia Martin, Shawn Christensen, Dino Sbardellati, Gillian Bergmann, and Jacob Francis for feedback throughout the project. Additionally, we thank L.G. Cardoso for their assistance with field work and laboratory experiments. We also thank Corwin Vannette for their assistance with floral nectar survey sample processing. We are grateful to the UC Davis Botanical Conservatory and UC Davis Arboretum staff as well as the UC Davis grounds crew for allowing us to conduct this work with plants on the UC Davis campus. This work is supported in part by an NSF REPS Postbaccalaureate Research Fellowship to LL and DEB # 1846266 to RLV. The authors declare no conflict of interest.

## Supporting Information

See supporting information for media recipes, additional methods on peroxide quantification and supplementary Figure S1.

