## Supporting Information for "Plant species vary in nectar peroxide, an antimicrobial defense tolerated by some bacteria and yeasts"

Table of Contents

Pg 2-3. Media recipes

Pg 4. Hydrogen Peroxide Quantification

Pg 5. Supplementary Figure S1

**Media Recipes**

Agar growth media:

Yeast media (YM) agar - *Metschnikowia reukaufii* and *Aureobasidium pullulans*

500 mL DI water
1.5 g malt extract
2.5 g peptone
5 g glucose
1.5 g yeast extract
10 g agar
After autoclaving, add 500 uL chloramphenicol in methanol (100 mg/mL)

Tryptic soy (TS) agar - *Neokomagataea thailandica* and *Acinetobacter pollinis*

500 mL DI water
7.5 g agar
2.5 g soytone
2.5 g sodium chlorine (NaCl)
25 g fructose
After autoclaving: 500 uL cycloheximide in methanol (100 mg/mL)

de Man, Rogosa and Sharpe (MRS) agar - *Apilactobacillus micheneri*

500 mL DI water
26 g MRS broth powder (Remel)
10 g fructose
7.5 g agar
After autoclaving: 500 uL cyclohexamide in methanol (100 mg/mL)

Liquid growth media:

Aim 3 artificial nectar, as described in (Mueller et al., 2023)

250 mL DI water
250 mL non-essential amino acids
37.5 g sucrose
18.75 g glucose
18.75 g fructose
5 g peptone

15 g yeast extract

Filter with 0.2 micron sterile filter

Aim 5 artificial nectar

500 mL DI water

40 g sucrose

80 g glucose

80 g fructose

4 g peptone

Filter with 0.2 micron sterile filter

<u>Experimental control solutions:</u>

Glucose oxidase 40% sugar substrate solution

250 mL DI water

80 g glucose

20 g sucrose

Filter with 0.2 micron sterile filter

**Hydrogen Peroxide Quantification**

Amplex™ Red Hydrogen Peroxide/Peroxidase Assay Kit (ThermoFisher, Waltham, MA):
In the presence of hydrogen peroxide, a peroxidase reacts with the colorless reagent, Amplex Red, to form a red oxidation product, the absorbance of which can be measured at 560 nm.

$$Amplex\ red\ +\ Horseradish\ Peroxidase\ +\ H_2O_2 \rightarrow Resorufin\ (red\ fluorescent\ oxidation\ product) + O_2$$

We performed this assay on 5 uL of floral nectar, and the absorbance was measured on a spectrophotometer microplate reader (model SYNERGY HTX). A standard curve was used to determine the absorbance to concentration conversion. Sample blanks (sample solution with no reagents added) and negative control samples lacking the addition of peroxidase were used to confirm lack of background absorbance and absence of natural peroxidases. Detection limit is 35 uM hydrogen peroxide because higher concentrations lead to sufficiently excess level of peroxide that oxidize resorufin back to the non-red Amplex Red form, resulting in artificially low peroxide levels. This requires sufficient serial dilution to determine the linear range of absorbance for a sample, which presents limitations when working with very small volumes of floral nectar.

Peroxide test strips (WaterWorks™ Mid Range Peroxide Check, Industrial Test Systems Inc., Rock Hill, SC):
0.2 - 5 uL of nectar was pipetted onto a test strip and the peroxide concentration was approximated visually with a color chart. We note that these test strips are non-specific and measure the concentration of all peroxides that may be present in a sample, rather than targeting solely hydrogen peroxide.

**Supplementary Figure S1:**

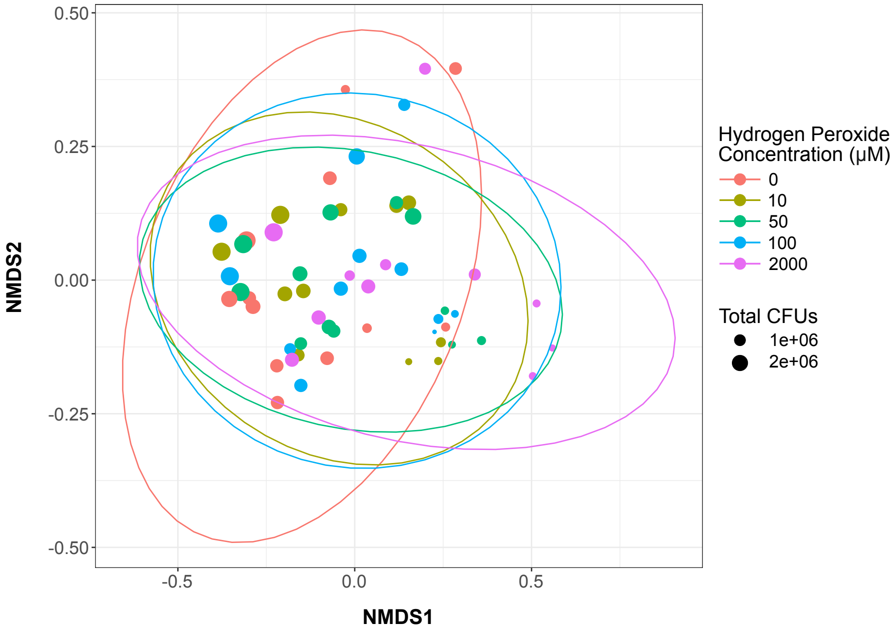


**Supplemental Figure S1:** NMDS of the microbial community samples to show impact of hydrogen peroxide concentration on microbial composition (PerMANOVA: $F_{4,58} = 0.75$, $p = 0.63$). Colors represent hydrogen peroxide concentration levels. The size of the filled circle illustrates the total microbial abundance. Ellipses represent 95% confidence intervals based on a multivariate T distribution.
